# Friend or foe: A mosquito parasite with mixed transmission mode displays mutualistic traits promoting oogenesis

**DOI:** 10.1101/2025.08.28.672828

**Authors:** Maxime Girard, Mathieu Laÿs, Edwige Martin, Laurent Vallon, An-nah Chanfi, Mélanie Bretton, Aurélien Vigneron, Séverine Balmand, Patricia Luis, Anne-Emmanuelle Hay, Claire Valiente Moro, Guillaume Minard

**Author notes:** Guillaume Minard. **Email:**. **Author Contributions:** Conceptualization: MG, AEH, CVM, GM; Experiments: MG, ML, MB, AC, EM, LV, SB; Bioinformatics and statistical analyses: MG, AV, GM; Supervision: AEH, CVM and GM; Writing – original draft: MG, AEH, CVM, GM; Writing – review & editing: All authors contributed. **Competing Interest Statement:** Authors declare they have no competing interests.

## Abstract

Mutualism is often selected in vertically transmitted symbionts due to their fitness interdependence with hosts. However, the evolution of mutualism remains unclear in symbionts using both vertical and horizontal transmission. In this study, we show that *Ascogregarina taiwanensis*, previously known as a weak horizontally transmitted parasite of the Asian tiger mosquito (*Aedes albopictus*), exhibits mutualistic traits that enhance mosquito reproduction. The symbiont improves embryogenesis and extends the egg-laying period while most females are pseudo-vertically transmitting symbiont oocysts to their progeny at oviposition sites. Dual transcriptomic analyses reveal that early oogenesis in infected females involves increased nitrogen metabolism in both partners, enhanced detoxification of blood waste, and activation of egg development pathways. These changes lead to improved assimilation of blood proteins essential for egg production. Our findings provide rare empirical evidence of a symbiont displaying both parasitic and mutualistic traits, offering new insights into the evolutionary dynamics of mixed-mode transmission symbioses.

**Significance Statement:** How mutualism evolves from parasitism remains a central question in evolutionary biology, particularly for symbionts that combine vertical and horizontal transmission. We show that *Ascogregarina taiwanensis*, an Apicomplexan parasite previously known to be costly for developing *Aedes albopictus* mosquitoes, is also pseudo-vertically transmitted: most females release oocysts into the water during oviposition, exposing their offspring. Surprisingly, infection enhances host reproduction by promoting oogenesis through protein assimilation, leading to the production of larger larvae. Dual transcriptomic analyses reveal coordinated shifts in host and parasite metabolism, especially in nitrogen assimilation. Our findings provide rare evidence that mutualistic traits can emerge in a symbiont with mixed-mode transmission, offering new insights into the evolutionary transitions shaping host–microbe interactions.

## Introduction

Host-microbe interactions are commonly classified along a continuum ranging from parasitism to mutualism, depending on their negative, neutral, or positive effects on each partner (1). While environmental conditions and genetic backgrounds modulate these interactions (2), the degree of dependency between partners also plays a crucial role in shaping the trajectory of the relationship (3). In particular, interdependency between partners is strongly influenced by the mode of transmission: vertical transmission tends to increase this dependency (4).

Many mutualists and parasites are thought to have evolved from free-living ancestors, some of which have transitioned into obligate symbionts over generations (5). Empirical evidence suggests that parasitism is often the initial step following a free-living lifestyle (5–7). For example, the free-living, photosynthetic ancestors of Apicomplexa lost chloroplast genes over the course of evolution and shifted from autotrophy to heterotrophy (8). To compensate for the loss of essential metabolic capabilities, Apicomplexa became highly dependent on the animal hosts they parasitize for nutrient acquisition. In some cases, interactions between hosts and environmental microbes became so critical that mutualistic relationships evolved. For instance, certain *Pantoea* species transitioned from free-living soil bacteria to obligate mutualists inhabiting specialized midgut crypts in the stinkbug *Plautia stali* (9). This association is maintained overtime through vertical transmission, *i.e*. females coat the egg masses with secretions containing symbionts, which are then ingested by the newly hatched nymphs (10).

Inter-individual transmission of microorganisms is a major force shaping the nature of host–microbe interactions (11). Vertical transmission, in particular, promotes tighter host–microbe associations and can lead to reduced virulence and the evolution of mutualistic traits that enhance the reproduction of both partners (5). In contrast, when symbionts are transmitted horizontally, their fitness becomes decoupled from that of the host, favoring the evolution of parasitic strategies (1, 12). For example, substantial evidence suggests that the aphid symbiont *Serratia symbiotica* was originally a horizontally transmitted parasite that evolved mutualistic characteristics under the selective pressure of vertical transmission (13, 14). Similarly, the virulence of the free-living alga *Symbiodinium microadriaticum* has been shown to decrease under experimental vertical transmission in jellyfish hosts (15). Together, empirical studies and controlled experiments demonstrate that vertical transmission tends to reduce microbial virulence and may promote the emergence of mutualism (16). However, direct observations of an ongoing transition from parasitism to mutualism driven by vertical transmission remain scarce in the literature.

The Asian tiger mosquito, *Aedes albopictus*, is the most invasive vector of vertebrate pathogens, and its global spread is a major concern for public health policies (17). Field populations of *Ae. albopictus* are commonly and heavily parasitized by *Ascogregarina taiwanensis*, a low-virulence Apicomplexan entomoparasite (18). This parasite negatively affects host survival and reproduction under nutrient-deficient conditions (19) or when infection loads are high (20). However, it has also been suggested that *As. taiwanensis* may have contributed to the competitive success of *Ae. albopictus* over other mosquito species during the invasion of new habitats—such as breeding sites also colonized by *Ae. aegypti* (21). This may be due to the higher virulence of *As. taiwanensis* toward *Ae. aegypti* larvae, while exerting limited impact on the fitness of its natural host (22). Consequently, *As. taiwanensis* may have either negative or indirectly beneficial effects on *Ae. albopictus*.

Infection by *As. taiwanensis* occurs when larvae ingest oocysts in breeding water, after which the parasite develops in the midgut and Malpighian tubules of pupae and adults (23). It is then disseminated by adult mosquitoes via a supposedly mixed transmission mode (24). Briefly, horizontal transmission occurs when emerging adults release meconium in aquatic habitats, or when infected individuals defecate or die in water. Several studies have also suggested a potential pseudo-vertical transmission via egg smearing from females to their offspring (20, 24), although this has yet to be clearly demonstrated.

In this study, we first investigated whether *As. taiwanensis* can be vertically transmitted from mother to offspring. We found that this transmission route is indeed possible, even though it co-occurs with horizontal transmission in breeding sites. Given the close link between female reproduction and parasite transmission, we then examined the impact of infection on mosquito oogenesis, fertility, fecundity, and offspring quality. We showed that, although parasitized and non-parasitized mosquitoes ingest similar amounts of blood, infection promotes oogenesis in females, resulting in larger eggs laid over an extended oviposition period and ultimately yielding larger larvae. Dual RNA-seq conducted across reproductive stages revealed that pathways involved in the assimilation of blood-derived nutrients—particularly nitrogen—were upregulated in both the parasite and the mosquito. In addition, parasitized females exhibited a more efficient turnover of protein digestion by-products during early oogenesis and greater protein assimilation toward the end of oogenesis.

## Results

### *As. taiwanensis* is pseudo-vertically transmitted through the aquatic habitat

To understand how *As. taiwanensis* persists across generations, we first examined its prevalence in females at different stages of the reproductive cycle. The proportion of females harboring oocysts remained high and stable in unmated (UNMAT; 81.2%), mated (MATED; 87.5%), and recently blood-fed females (86.7%; 1 day after blood meal, 1DABM) (**Fig. 1a**; **Table 1**). However, oocyst prevalence sharply dropped during late oogenesis (3DABM) to an average of 40%, suggesting parasite release during oviposition (**Fig. 1b**).

**Table 1.**
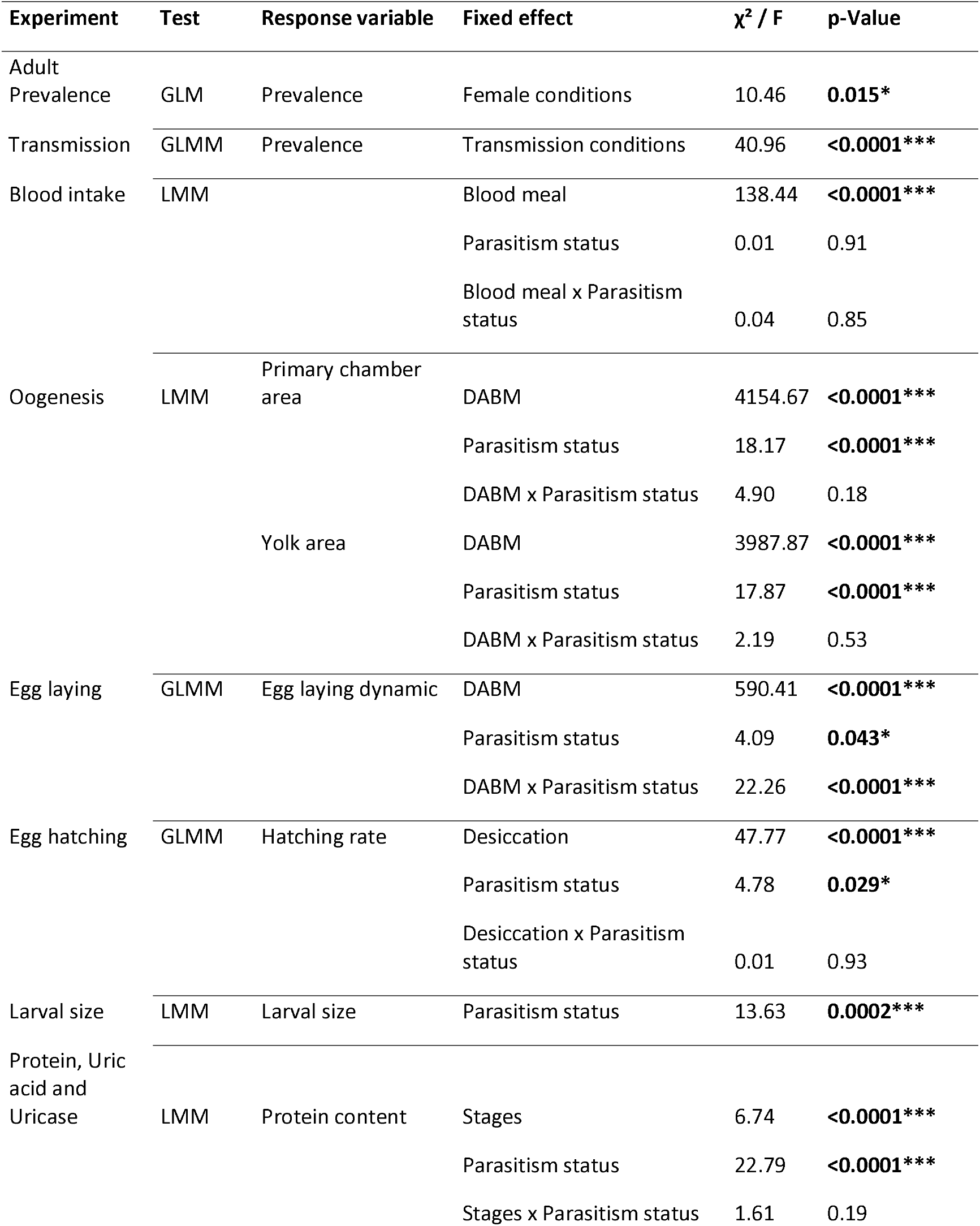
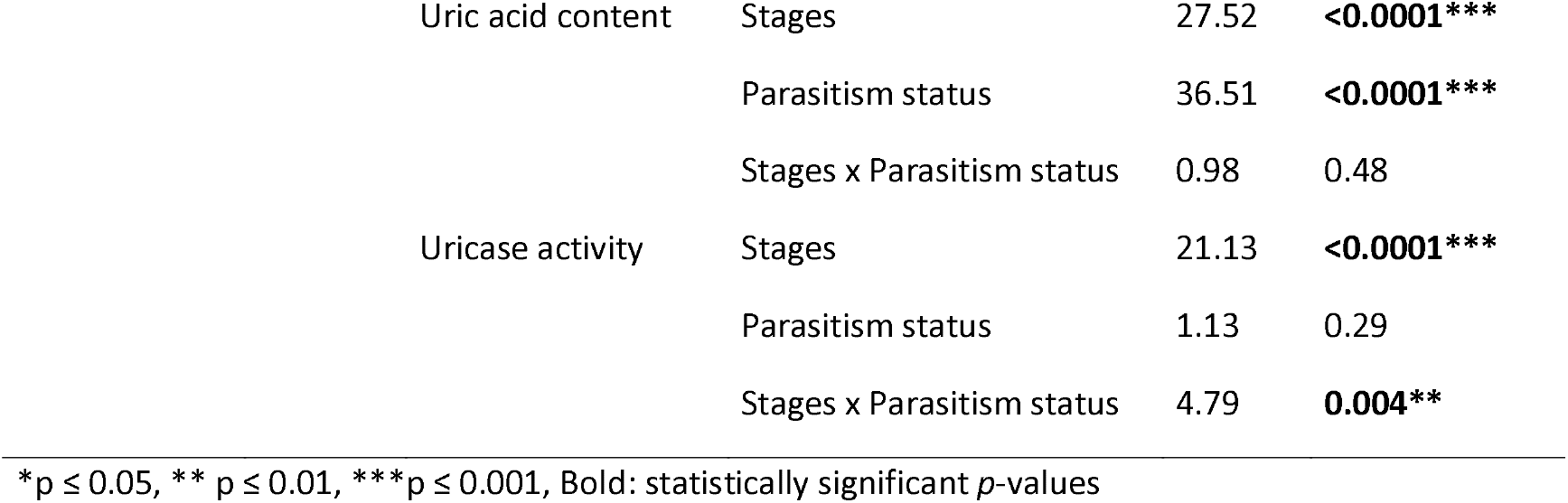
Statistical analysis summary.

**Figure 1.**
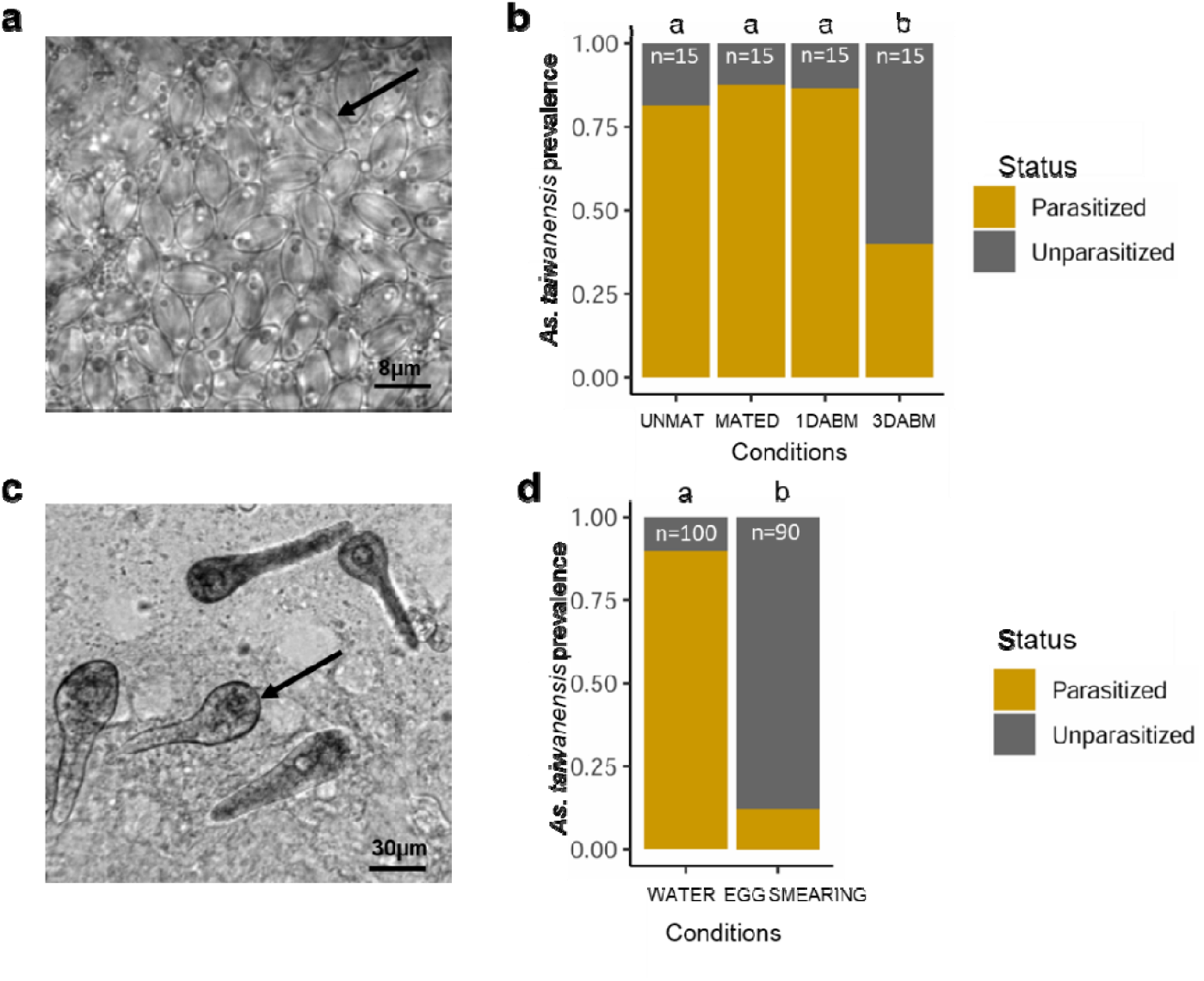
Prevalence of *As. taiwanensis* in females over life stages and vertical transmission. (a) The prevalence of *As. taiwanensis* oocysts was measured in females. The image was acquired using light microscopy at ×1000 magnification; the black arrow indicates a single oocyst. (b) Prevalence of oocysts in female Malpighian tubules at different physiological stages: unmated (UNMAT), mated (MATED), one day after blood meal (1DABM), and three days after blood meal (3DABM). (c) The prevalence of *As. taiwanensis* was estimated in larvae in its trophozoïte. The image was acquired using light microscopy at ×400 magnification; the black arrow indicates a single trophozoite. (d) Vertical transmission assessed in newly hatched larvae exposed either to water from oviposition cups (WATER) or to sterile water after egg surface sterilization (EGG SMEARING). Compact letter displays indicate statistically significant differences based on post hoc Tukey HSD tests.

To test whether females transmit the parasite to their offspring via the aquatic habitat, we measured the prevalence of *As. taiwanensis* trophozoites in larvae. When larvae developed in the same water in which their parasitized mothers laid eggs, infection prevalence reached 89%. In contrast, only 12.2% of larvae were infected when eggs were transferred to sterile water before hatching (**Fig. 1c, 1d**; **Table 1**). These results indicate that pseudo-vertical transmission occurs predominantly via oocysts released in water, rather than through egg smearing.

### Parasitized females produce larger eggs over an extended laying period and give rise to larger larvae

As. taiwanensis colonizes the Malpighian tubules—organs involved in excretion and osmoregulation, particularly active during blood meal digestion. Following blood feeding, both parasitized and unparasitized females ingested similar blood volumes, as inferred from comparable abdomen width (Fig. 2a; Table 1).

**Figure 2.**
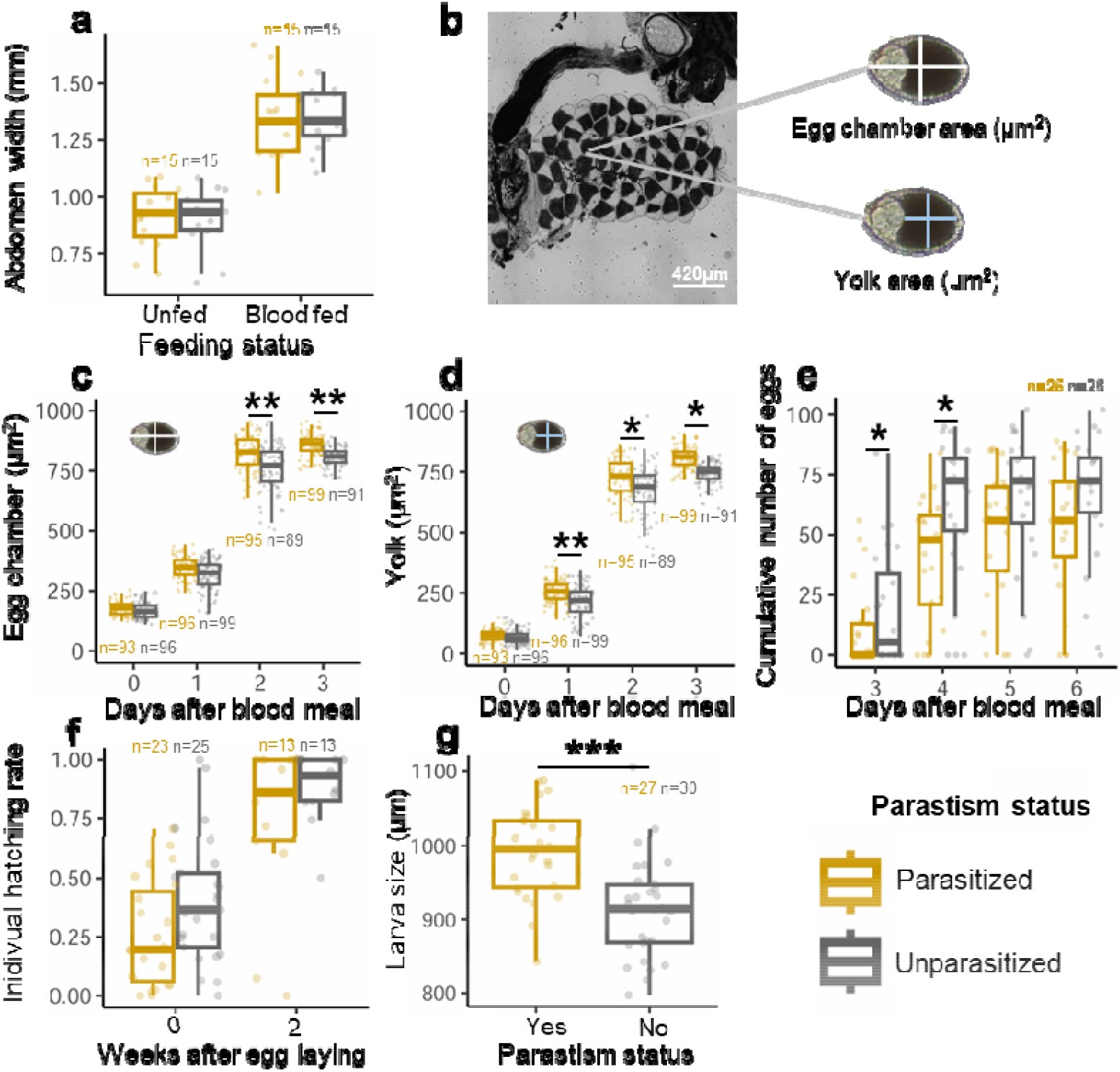
Impact of *As. taiwanensis* on female blood feeding, oogenesis, egg laying, egg hatching and larval size. (a) Abdomen width was measured in parasitized and unparasitized females, before (Unfed) and after a blood meal (Blood fed). (b) Representative image of primary follicle chamber and yolk area in a parasitized female one day after blood feeding (1 DABM), visualized using light microscopy (×100 magnification). (c) Area of primary follicle chambers and (d) yolks was quantified daily from 0 to 3 DABM in parasitized and unparasitized females. (e) Egg-laying dynamics were monitored daily from 3 to 6 DABM in parasitized and unparasitized females and represented as cumulative egg numbers. (f) Hatching success was measured with (2 weeks) and without (0 weeks) a desiccation period in both parasitized and unparasitized groups. (g) Larval size was assessed after hatching for parasitized and unparasitized individuals. Asterisks indicate significant differences (^*^p ≤ 0.05, ^**^p ≤ 0.01, ^***^p ≤ 0.001; Tukey HSD post hoc test).

Despite similar blood intake, parasitized females showed enhanced oogenesis. Primary egg chamber area increased by 7.8% at 2DABM (t=3.31, p=0.001^**^) and 6.5% at 3DABM (t=2.94, p=0.004^**^), while yolk (vitellus) area increased by 19.8% at 1DABM (t=2.66, p=0.001^**^), 8.1% at 2DABM (t=2.03, p=0.02^*^), and 8.6% at 3DABM (t=2.64, p=0.01^*^) (Fig. 2b–d).

Egg-laying was temporally shifted in parasitized females: a larger fraction of their eggs were laid at 4DABM (72.6%), compared to 93.0% at 3DABM in unparasitized females (Fig. 2e). This delay was significant at both 3 and 4DABM (z=-2.23, p=0.026^*^; z=-2.14, p=0.032^*^). However, the total number of eggs laid was similar across groups (z=-1.46, p=0.144), as was hatching success, whether eggs were fresh or stored (Fig. 2f).

Importantly, larvae hatching from eggs laid by parasitized females were significantly larger (+8.4%) than those from unparasitized females (Fig. 2g; Table 1), suggesting a developmental advantage potentially linked to improved maternal resource allocation.

### Parasitism and reproductive stage shape the mosquito transcriptome

We next examined transcriptomic changes in female mosquitoes across reproductive stages and infection status (UNMAT, MATED, 1DABM, and 3DABM). Of 36,525 transcripts, 30,827 were retained for downstream analysis. PCA revealed that both female life stage and parasitism status significantly structured gene expression profiles (Fig. 3a), accounting for 49.6% of total variance: 34% by life stage (F=7.20, p=0.001^***^), 3.8% by infection status (F=2.39, p=0.067), and 11.8% by their interaction (F=2.51, p=0.012^*^).

**Figure 3.**
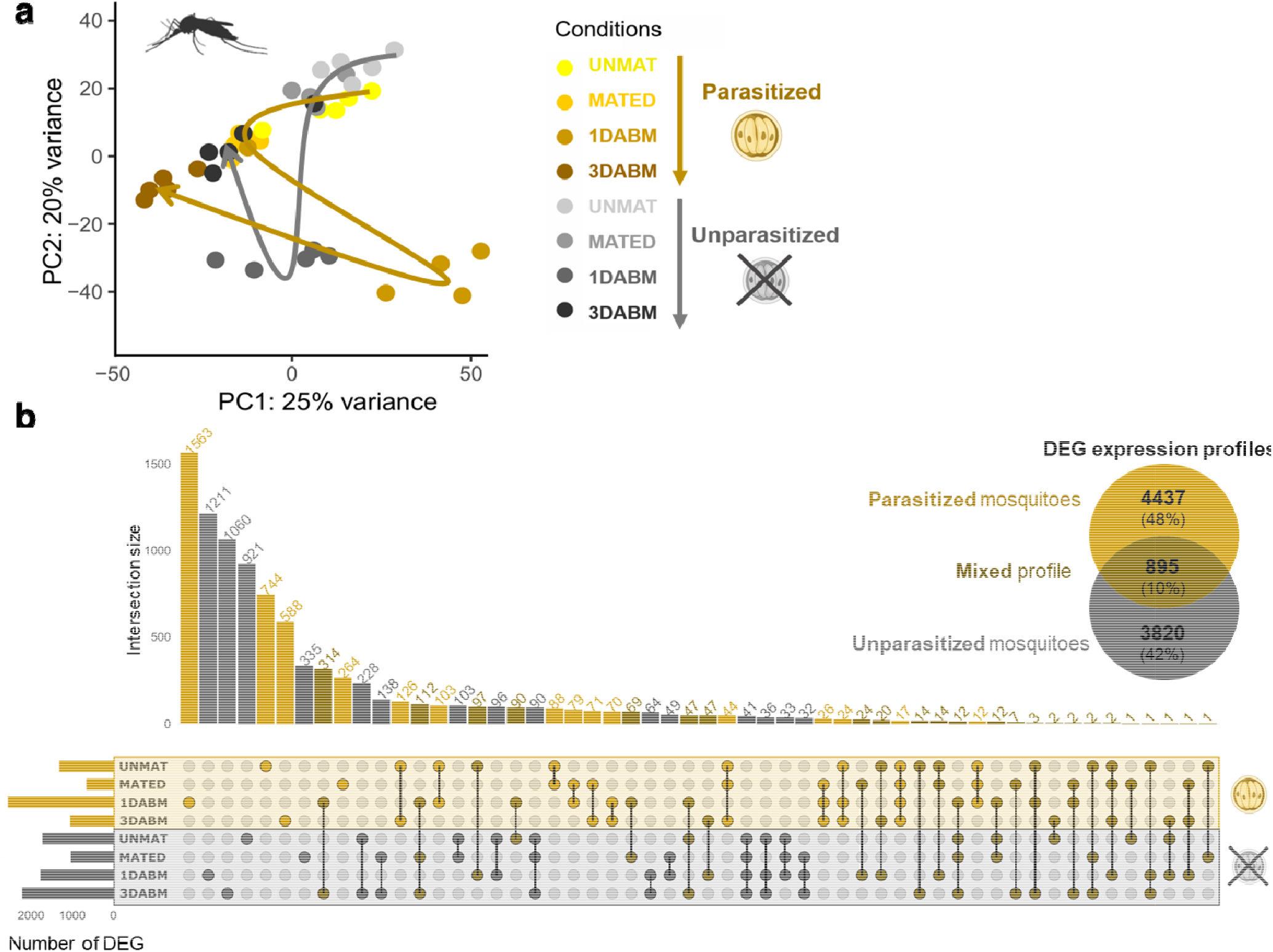
Transcriptomic impact of *Ascogregarina taiwanensis* on *Ae. albopictus* females: Principal Component Analysis and differentially expressed genes (DEGs). (a) Principal Component Analysis (PCA) of transcriptome profiles from 5 parasitized and 5 unparasitized females at four physiological stages: unmated (UNMAT), mated (MATED), one day after blood meal (1DABM), and three days after blood meal (3DABM). Each dot represents the transcriptome of an individual female. (b) A Venn diagram and an UpSet plot summarize the number and overlap of DEGs between parasitized and unparasitized mosquitoes across female conditions. A total of 9,152 DEGs were identified using pairwise comparisons and Wald tests with Benjamini-Hochberg correction. Intersections are shown as columns. Colors indicate genes specific to parasitized mosquitoes (yellow), unparasitized mosquitoes (grey), or shared between both conditions (dark yellow). Female physiological stages are color-coded and labeled as UNMAT, MATED, 1DABM, and 3DABM.

Differentially expressed genes (DEGs) between parasitized and unparasitized mosquitoes were analyzed (**Fig. 3b**). Each DEG represents a transcript for which the expression abundance is significantly higher either in parasitized or unparasitized mosquitoes. In total, 9,152 differentially expressed genes (DEGs) were identified (Fig. 3b), with 4,437 and 3,820 genes specific to parasitized and unparasitized females, respectively, and 895 shared. The greatest divergence was observed at 1DABM, with 2,626 DEGs, including 1,563 specific to parasitized females.

### Parasitized mosquitoes upregulate genes related to development and protein metabolism after blood feeding

GO enrichment analysis showed that parasitism drives distinct gene expression programs depending on reproductive stage (**Fig. 4**). In UNMAT and MATED females, parasitized individuals overexpressed genes involved in macromolecule, lipid, nucleoside, and organic acid metabolism. Enrichment of GO terms associated with lipid (*e.g*. fatty acids) and nucleoside (*e.g*. UDP-glucose metabolism) processing was more pronounced in MATED parasitized individuals (52% and 17% of the total number of genes respectively). Conversely, transcriptome profiles of unparasitized mosquitoes were similar during the UNMAT and MATED life stages. They upregulated functions related to nitrogen (*e.g*. nitrogen/protein metabolism, amide and nitrogen compound biosynthesis), macromolecule (*e.g*. biosynthesis of compounds such as purine) metabolism as well as mitochondrial (*e.g*. ATP synthesis, anion, electron and ATP transport) activity.

**Figure 4.**
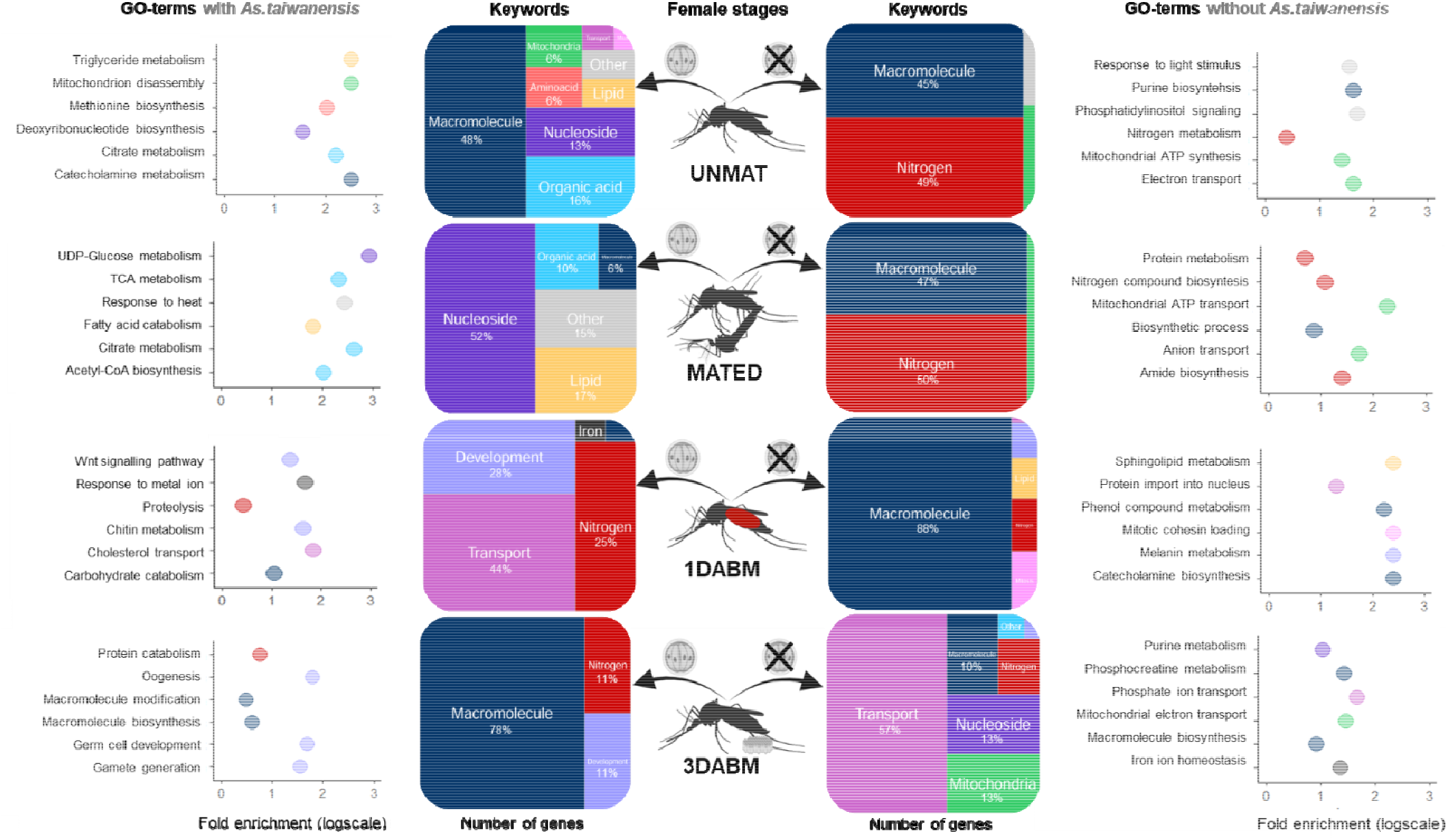
Functional annotation of differentially expressed genes (DEGs) between parasitized and unparasitized*Ae. albopictus* females across life stages. Treemaps display enriched functional categories summarizing Gene Ontology (GO) terms associated with DEGs at four female physiological stages: unmated (UNMAT), mated (MATED), one day after blood meal (1DABM), and three days after blood meal (3DABM). The surface area of each keyword reflects the proportion of associated DEGs. For each stage and infection status, the fold enrichment scores of the top six GO terms are shown (log scale). Percentages of genes involved in each functional category are displayed when exceeding 1%.

After blood feeding, transcriptomic differences intensified. At 1DABM and 3DABM, parasitized females overexpressed genes involved in development (e.g. wnt signaling pathway, chitin metabolism, oogenesis germ cell development and gamete generation) and nitrogen metabolism (e.g. proteolysis/protein catabolism; **Fig. 4**). In contrast, DEGs in 1DABM unparasitized females were dominated by macromolecule metabolism (88%).

Iron metabolism was also more active in parasitized 1DABM females (2% of DEGs) than in 3DABM unparasitized females (<1%). A focused analysis of enriched metabolic pathways in 1DABM parasitized females (Table S1) revealed upregulation of amino acid metabolism, detoxification (glutathione, cytochrome P450), lipid catabolism, hormonal regulation, vitamin metabolism (ascorbate) and energy (pyruvate) pathways.

Notably, several serine protease-like genes involved in protein catabolism showed very high fold changes (129.7, 55.4, 38.6). Genes for glutamine synthetase and glutamate dehydrogenase-like enzymes (GS/GOGAT cycle) were also highly upregulated (fold changes: 6.9, 6.1), as were detoxification genes like glutathione S-transferase-like (fold change: 116.5) and peroxiredoxin (8.1). Hydroxysteroid 17-beta dehydrogenase 11 (HSD17B11), involved in lipid metabolism and hormonal regulation, was upregulated by 20.8-fold.

### The transcriptome of *As. taiwanensis* remains mostly stable across mosquito life stages

Given the strong effect of parasitism on the mosquito transcriptome, we also examined the transcriptional dynamics of the symbiont *As. taiwanensis* across female life stages. PCA conducted on the parasite gene expression profiles (based on 1,679 transcripts retained after abundance estimation and normalization from 4,676 identified) revealed no significant global variation among life stages (F = 1.10, p = 0.149; **Fig. 5a**). This suggests that the overall *As. taiwanensis* transcriptome remains relatively stable across conditions.

**Figure 5.**
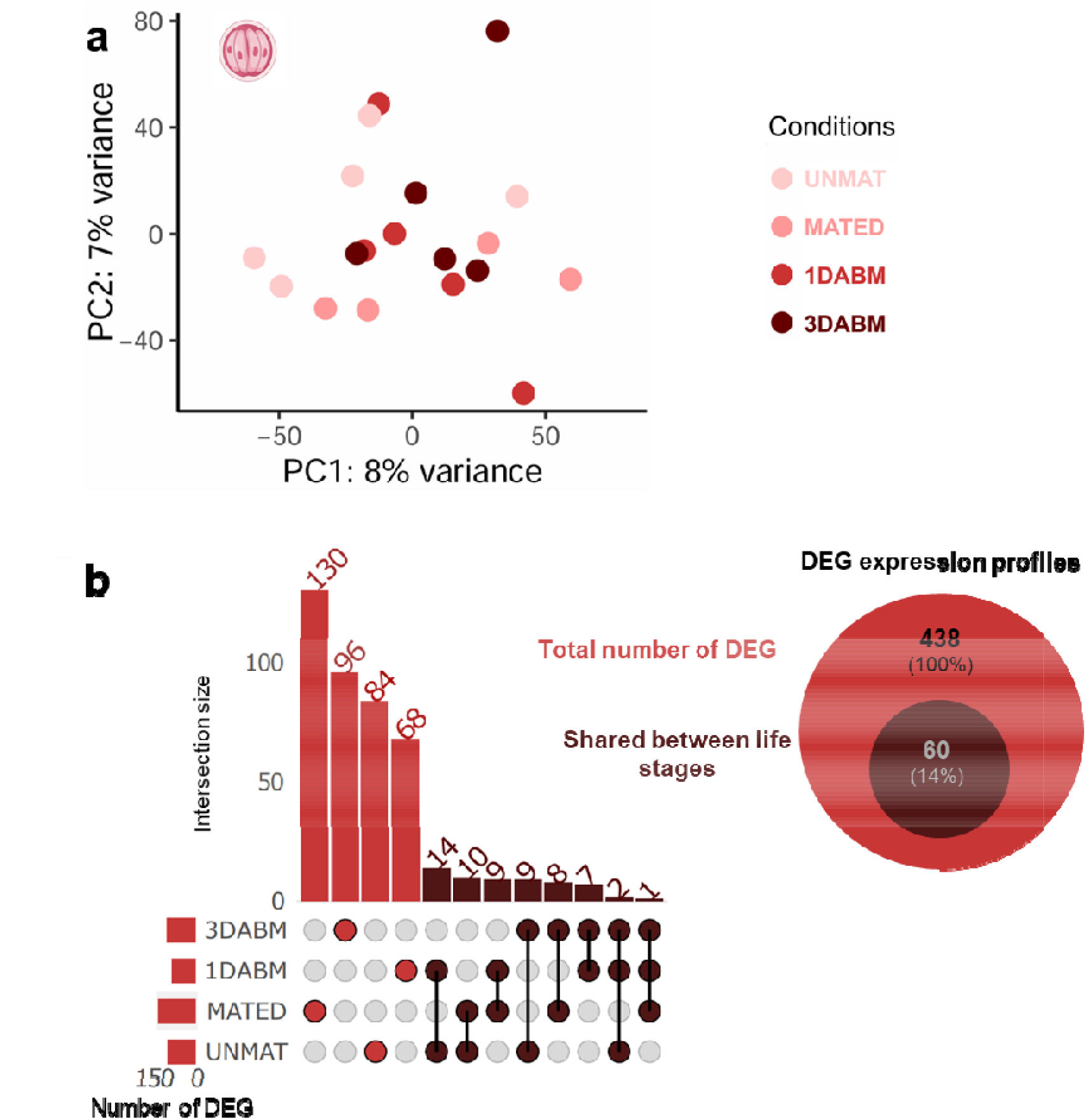
Principal Component Analysis and differentially expressed genes (DEGs) in the *Ascogregarina taiwanensis* transcriptome. (a) Principal Component Analysis (PCA) of de novo transcriptome profiles from *As. taiwanensis* within 5 parasitized *Ae. albopictus* females across four physiological stages: unmated (UNMAT), mated (MATED), one day after blood meal (1DABM), and three days after blood meal (3DABM). Each dot represents the parasite transcriptome associated with an individual host. (b) A Venn diagram and UpSet plot summarize the number and overlap of DEGs identified across stages. A total of 438 DEGs were detected using pairwise comparisons and Wald tests with Benjamini-Hochberg correction. Among them, 378 DEGs were specific to a single life stage (red), while 60 were shared across multiple stages (dark red).

Despite the absence of a major global pattern, differential gene expression (DGE) analysis identified 438 DEGs across life stages. Among them, 60 genes were shared between conditions, while stage-specific DEGs ranged from 68 to 130 (**Fig. 5b**).

### Transcriptomic variation in *As. taiwanensis* during mosquito reproduction

To provide a better overview of the mechanisms involved in host-parasite interactions, functional annotations of the *As. taiwanensis* DEG were also performed (**Fig. 6**). The main GO term keywords associated with genes expressed in *As. taiwanensis* were related to macromolecule, nitrogen and nucleoside metabolism. Nitrogen metabolism, which was particularly abundant during the early phase of mosquito blood digestion and oogenesis (*i.e*. 1DABM with 28% of the genes), was consistently the second most important keyword across female life stages. This includes genes involved in protein metabolism, particularly glutamine metabolism (**Fig. 6**). For instance, the *glutamine amidotransferase* like, which facilitates the extraction of ammonia from glutamine, was identified. The arginine biosynthesis pathway was also notably enriched in both 1DABM and 3DABM conditions (**Table S2**). In contrast, the C5-branched dibasic acid metabolic acid pathway, which contributes to the synthesis of precursors for branched-chain amino acids such as leucine, isoleucine and valine, was only enriched in 1DABM. Additionally, genes related to macromolecule GO-terms were more abundant in blood-fed females (1DABM and 3DABM), especially those involved in the tricarboxylic acid cycle.

**Figure 6.**
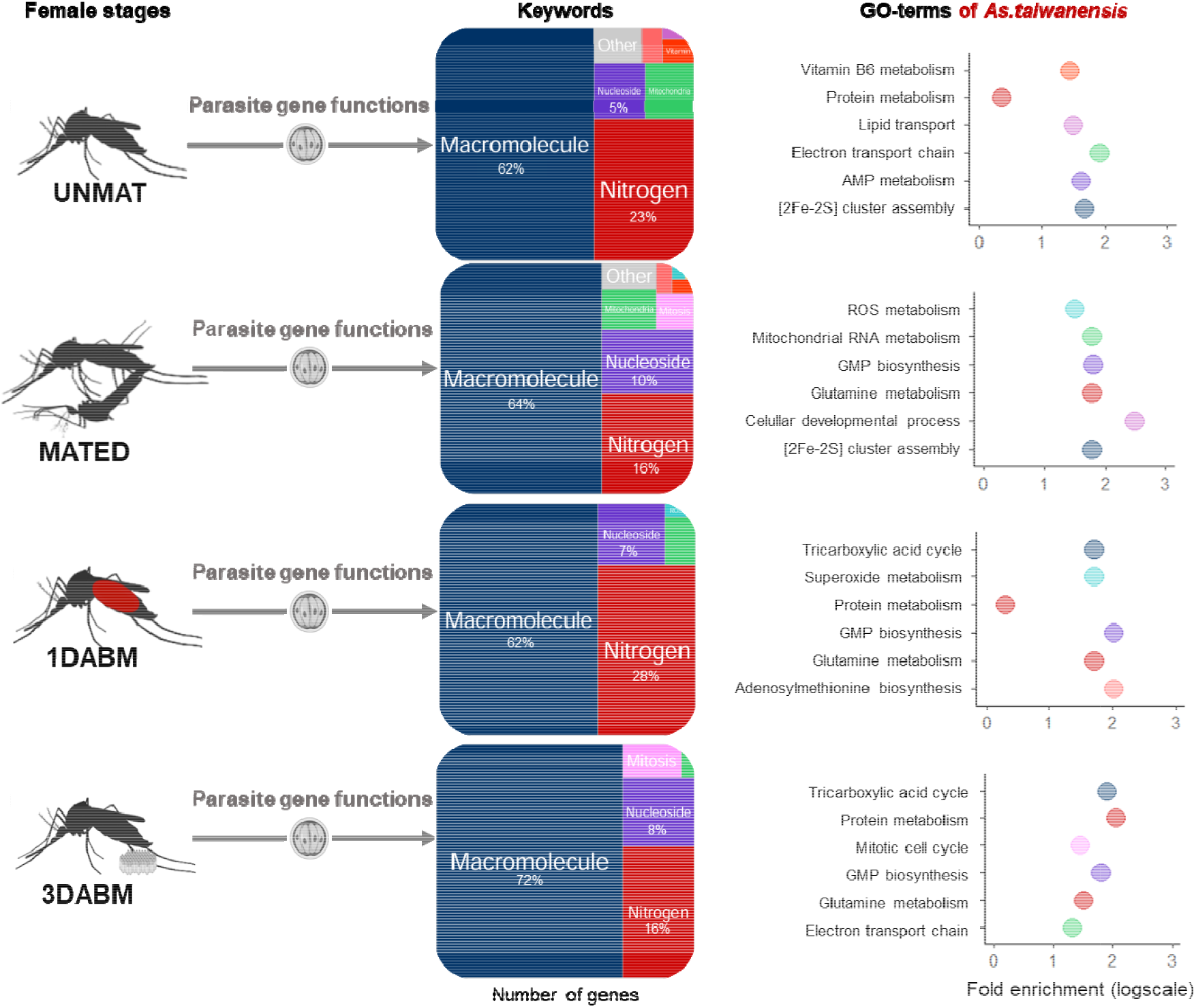
Functional annotation of *Ascogregarina taiwanensis* transcriptome across mosquito female life stages. The *As. taiwanensis* transcriptome associated with *Ae. albopictus* females was functionally annotated at four physiological stages: unmated (UNMAT), mated (MATED), one day after blood meal (1DABM), and three days after blood meal (3DABM). GO terms were simplified into functional keywords and visualized as treemaps. The surface area of each keyword reflects the proportion of DEGs it aggregates (log scale). For each life stage, the top six enriched GO terms are shown with their fold enrichment scores (logarithmic scale). Percentages of genes involved in each keyword are displayed when exceeding 1%.

To further explore potential stage-specific functional shifts in *As. taiwanensis*, we annotated the identified DEGs (**Fig. 6**). The most enriched GO terms were related to macromolecule metabolism, nitrogen metabolism, and nucleoside metabolism. Nitrogen metabolism was particularly prominent at 1 day after blood meal (1DABM), representing 28% of DEGs at this stage. This included genes involved in protein and glutamine metabolism, such as a *glutamine amidotransferase*-like gene, which facilitates ammonia extraction from glutamine. Additionally, the arginine biosynthesis pathway was significantly enriched in both 1DABM and 3DABM (**Table S2**). In contrast, the C5-branched dibasic acid metabolism pathway—implicated in the synthesis of branched-chain amino acid precursors (e.g., leucine, isoleucine, valine)—was specifically enriched at 1DABM.

Genes associated with macromolecule-related GO terms, particularly those involved in the tricarboxylic acid (TCA) cycle, were more abundant in blood-fed females (1DABM and 3DABM), suggesting a metabolic shift aligned with the host’s reproductive physiology.

### A potential mutualistic role of *As. taiwanensis* in blood resource assimilation

To test whether *As. taiwanensis* contributes to enhanced resource assimilation during oogenesis, we conducted complementary experiments. Under blood dilution treatments, parasitized females were less impacted than their unparasitized counterparts. When fed blood diluted fivefold, unparasitized females failed to produce eggs, while parasitized females formed an average of 10 ± 7 eggs (**Fig. S1**). These results suggest that *As. taiwanensis* may enhance nutrient assimilation, particularly of proteins, during reproduction.

To investigate this further, we measured total protein content in females across reproductive stages. In both groups, protein levels increased by ∼50% at 1DABM and declined by 3DABM. However, parasitized females showed significantly higher protein levels at 3DABM, accumulating 37.8% more protein than unparasitized ones (**Fig. 7a**).

**Figure 7.**
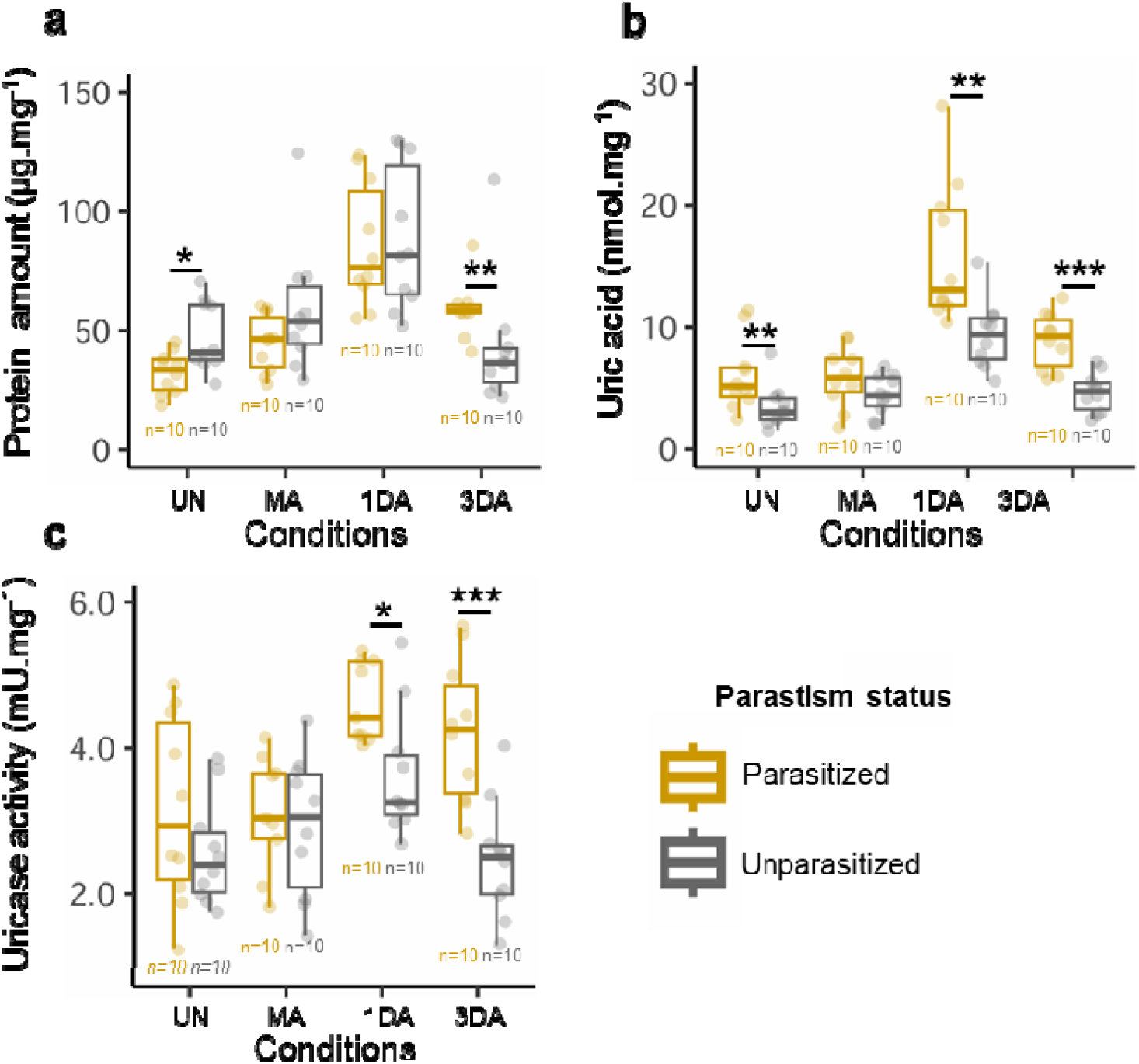
Quantification of proteins, uric acid, and uricase activity in parasitized and unparasitized *Ae. albopictus* females across physiological stages. (a) Total protein content, (b) uric acid concentration, and (c) uricase activity were measured in parasitized and unparasitized females at four life stages: unmated (UN), mated (MA), one day after blood meal (1DA), and three days after blood meal (3DA). Measurements were performed on 10 biological replicates per condition, each consisting of three pooled females and normalized to dry mass. Asterisks indicate statistically significant differences (^*^p ≤ 0.05, ^**^p ≤ 0.01, ^***^p ≤ 0.001; Tukey HSD post hoc test).

We then quantified uric acid levels and uricase activity—two indicators of nitrogenous waste metabolism (25)—to assess whether increased protein levels in parasitized females reflected more efficient protein digestion. Both uric acid (**Fig. 7b**) and uricase activity (**Fig. 7c**) increased significantly between 1DABM and 3DABM, particularly in parasitized females. These findings support the hypothesis that *As. taiwanensis* enhances protein assimilation and nitrogen metabolism during blood digestion and contribute to oogenesis.

## Discussion

Empirical studies have shown that vertical transmission favors the evolution of low-virulence or even beneficial parasitic traits (1). Mixed transmission modes—combining vertical and horizontal pathways—may also select for reduced virulence, depending on the relative importance of each route for host and parasite fitness (1, 5, 11). The pseudo-vertical transmission of *Ascogregarina* species, however, remains poorly characterized (24). Previous work suggested that males primarily mediate horizontal transmission by contaminating breeding water, while females disperse oocysts to new sites, mimicking vertical transmission (26). Since *As. taiwanensis* is expelled from adult mosquitoes without replicating in the adult stage (23), our results support the hypothesis that fewer parasites are released by females before oviposition, increasing the likelihood of transmission to their own offspring. This combination of dispersal and timing suggests a mixed transmission mode that may stabilize host–parasite association and promote the evolution of mutualistic traits.

Indeed, our findings indicate that *As. taiwanensis* positively affects mosquito reproduction: parasitized females produce larger progeny and sustain oogenesis even under limited blood resources. These benefits appear to be linked to enhanced nutrient assimilation, especially of nitrogenous compounds derived from blood proteins. Additionally, parasitized females exhibited altered oviposition behavior, distributing their eggs over a longer period than unparasitized females—a trait that could increase both reproductive success and parasite transmission.

Vertical transmission is often tightly coupled with host reproductive physiology, particularly when symbionts rely on host cellular machinery for follicle entry (27). For instance, *Wolbachia* and *Babesia bovis* hijack vitellogenin receptors to access ovarian tissues in true bugs (28) and ticks (29), respectively. Some symbionts even enhance host resource allocation to oocytes, as seen in *Nasuia deltocephalinicola*, which increases protein transport by 20% in the rice leafhopper *Nephotettix cincticeps* (30). While most such interactions involve direct contact with the ovary, *As. taiwanensis* remains confined to the Malpighian tubules and does not colonize ovarian tissues. Its release alongside eggs suggests a pseudo-vertical transmission route more similar to that observed in the bacterial symbionts *Pantoea* sp. of the green stink bug *Nezara viridula* (31, 32) or *Burkholderia* sp. in the oriental chinch bug *Cavelerius saccharivorus* (33). These facultative symbionts, transmitted through egg smearing or environmental contamination, also confer fitness benefits despite lacking direct contact with reproductive organs.

Our study further supports a functional mutualism between *As. taiwanensis* and *Ae. albopictus*, particularly during blood digestion and oogenesis. In mosquitoes, blood meal digestion triggers complex endocrine responses involving the brain, fat body, midgut, and ovaries (34). Proteins from the blood are broken down into amino acids that fuel yolk protein synthesis via the TOR pathway and promote follicle development through hormonal cascades involving insulin-like peptides and ecdysteroids (35). Concurrently, nitrogenous waste generated from protein catabolism—uric acid, urea, or ammonia—must be either excreted or detoxified (36) via the uricotelic or trans-deamination pathways (37), with amino acids like glutamine and proline acting as key intermediates (38).

In our experiments, parasitized females upregulated genes involved in proteolysis and protein catabolism, suggesting increased amino acid release. Transcriptomic signatures also pointed to enhanced oogenesis hormonal signaling, including overexpression of *HSD17B11-like*, part of the steroid biosynthesis pathway. Notably, the *GS-GOGAT* enzyme complex—central to ammonia assimilation into glutamine and glutamate—was also overexpressed in parasitized females (39), supporting improved nitrogen recycling.

Although *As. taiwanensis* is not known to be metabolically active in the adult mosquito, our transcriptomic analysis revealed that it remains transcriptionally active. Genes involved in nitrogen metabolism, particularly glutamine metabolism, were highly expressed. These findings raise the possibility that the parasite utilizes excess glutamine produced by the host, possibly via its own glutamine amidotransferase. Such reciprocal use of nitrogenous compounds suggests a metabolic interplay that benefits both partners through enhanced nitrogen turnover.

Symbiont-mediated nitrogen metabolism is well documented in phytophagous insects. Nitrogen-fixing or recycling microbes, such as diazotrophs in the long-horned beetle (40), *Buchnera aphidicola* in aphids (41), and *Ischyrobacter davidsoniae* in turtle ants (42), enable hosts to cope with nitrogen-poor diets. While mosquitoes ingest protein-rich blood (60–80 mg/mL) (43), they assimilate only a fraction of it: about 10% for oogenesis, 20% for storage, and the remainder is excreted as waste (44). Our results suggest that *As. taiwanensis* enhances protein assimilation and nitrogen detoxification, shifting this balance toward increased reproductive output.

Beyond oogenesis, parasitized females laid eggs over a longer period and produced larger larvae—two traits known to enhance offspring survival. Extended oviposition period may enhance “skip oviposition” behavior, a bet-hedging strategy in *Ae. albopictus* where eggs are spread across multiple breeding sites (45). Larger larvae generally outperform smaller ones in both intra- and interspecific competition, improving juvenile survival under natural conditions (46– 48). These offspring-level benefits also benefit the parasite, whose life cycle depends on successful host development and dispersal (23). Additionally, prolonged oviposition could expand the spatial reach of the parasite, increasing opportunities for horizontal transmission to unrelated individuals.

While our data suggest a mutualistic interaction, we cannot exclude the possibility of host compensatory plasticity. In many species, females increase investment in reproduction when facing threats like parasitism or predation, potentially as an adaptive response to reduce offspring mortality (49–52). Although we did not observe higher fecundity in parasitized females under normal conditions, differences in oviposition behavior and progeny size could still represent a plastic adjustment to infection. Notably, fecundity impairment occurred only in unparasitized females fed with diluted blood—conditions mimicking host defensive behavior and partial feeding in nature (53).

In summary, our study highlights the complex and potentially mutualistic nature of the interaction between *Ae. albopictus* and *As. taiwanensis*. The parasite is efficiently pseudo-vertically transmitted and may enhance mosquito reproductive success through improved blood digestion and nitrogen assimilation. Future studies should focus on tracking nutrient fluxes between host and parasite, particularly toward the ovaries, to further unravel the metabolic interdependence shaping this unusual interaction.

## Materials and Methods

### Mosquito lines

Parasitized and unparasitized *Aedes albopictus* lines originated from a hybrid laboratory population derived from two French natural populations sampled in Villeurbanne (N: 45°46′18.990″, E: 4°53′24.615″) and Pierre-Bénite (N: 45°42′11.534″, E: 4°49′28.743″) in 2017, following a previously described protocol (18). Both lines were reared under identical conditions in a biosafety level 2 insectary: 28°C, 80% relative humidity, and an 18h:6h light/dark cycle. Larvae were reared in 1.5 L of dechlorinated water and fed daily with a 25:75 yeast:fish food mixture (Biover; Tetra). Adults were kept in cages and provided with 10% sucrose *ad libitum. As. taiwanensis* presence was monitored at each generation by examining crushed pools of 25 adults and their rearing water under ×400 magnification using an optical microscope (Leica).

### Prevalence of *As. taiwanensis* across the female reproductive cycle

To track parasite prevalence throughout the female reproductive cycle, oocyst presence was recorded at key stages (**Figure S1**): unmated (UNMAT, 7 days old), mated (MATED, 14 days), 1 day after blood feeding (1DABF, 15 days), and 3 days after blood feeding (3DABF, 17 days). These timepoints correspond to pre-reproductive, host-seeking, early oogenesis, and post-oogenesis stages, respectively. Fifteen females per condition were individually crushed in 1.5 mL tubes with 100 µL sterile water and three 3 mm glass beads using a FastPrep lysis system (MP Biomedicals; 30 s, 20 m/s). Samples were observed at ×400 magnification to detect oocysts.

### Pseudo-vertical transmission assays

To assess pseudo-vertical transmission, blood-fed females were isolated 2 days post-blood meal in 50 mL tubes with 25 mL sterile water and a piece of blotting paper. After 3 days (end of oviposition), eggs were either kept in the same container (enabling waterborne and egg-smearing transmission) or transferred to a new container with sterile water (enabling only egg-smearing transmission). One milligram of food was added to each tube. Eggs were vacuum-incubated at −20 Hg for 6 h at 28°C. Five days post-hatching, third- and fourth-instar larvae were dissected under a stereomicroscope to detect *As. taiwanensis* trophozoites. Ten larvae per female were examined: 10 females for waterborne (n = 100 larvae), 9 for egg-smearing transmission (n = 90 larvae).

### Blood intake assessment

Unfed and blood-fed females were anesthetized on ice immediately after feeding. Abdomens were photographed at ×40 magnification, and width was measured in ImageJ v2.0 (54) for 15 individuals per condition. Abdomen width served as a proxy for blood intake (55).

### Oogenesis dynamics

To enhance vitellogenesis, 14-day-old females were starved for 6 h, then fed for 20 min using an artificial feeder (Hemotek) containing undiluted or diluted (1:2 or 1:5 with PBS) rat blood. The feeder was covered with pork intestine and maintained at 37°C. Females were dissected immediately (0 h), or 24 h, 48 h, or 72 h post-feeding. For each female, 6–10 primary follicles were photographed at ×100 magnification, and follicle and yolk areas were measured in ImageJ v2.0 (54). Experiments with diluted blood were performed on 5 females per condition (10 ovaries/female).

### Egg-laying dynamics, hatching rate, and larval size

Two days post-blood meal, 25 parasitized and 25 unparasitized females were individually placed in 50 mL tubes with 25mL of dechlorinated water and blotting paper. Females were transferred daily to new tubes for 6 days, and egg counts were recorded. Eggs were either immediately hatched or stored for 2 weeks on a dry blotting paper at 28°C and 80% relative humidity. To enhance hatching, they were vacuum-incubated (−20 Hg, 8 h, 28°C) with 1 mg fish food to stimulate hatching. For 13 parasitized and 14 unparasitized females, 5–8 first-instar larvae were photographed (×40), and size was measured in ImageJ v.2. (54). Total larval counts were recorded after one week.

### RNA extraction, library preparation and sequencing

Five parasitized and five unparasitized females were collected from each reproductive condition (*i.e*. UNMAT, MATED, 1DABM, 3DABM). Prior to euthanasia in liquid nitrogen, individuals were gently stimulated for 10 min to mimic natural activity. RNA was extracted following a published method (56). Libraries were prepared using the TruSeq Stranded mRNA kit (Illumina) and sequenced (2×150 bp) on a NovaSeq 6000 (Microsynth AG). Reads (17.7–30.8 million/sample) were demultiplexed and adapter-trimmed. Data are available at ENA (ERS23447863 will be available upon acceptance).

### Transcriptome analysis of *Ae. albopictus*

Reads were processed on Galaxy (https://usegalaxy.fr/) following Batut et al. (56). Quality was assessed with FastQC v0.12.1 (57), trimmed with Cutadapt v4.8 (58) with a minimum length of 75bp and a minimal quality score of 30, and mapped to the *Ae. albopictus* genome (RefSeq GCF_006496715.1) using STAR v2.7.11a (59). Mapped reads were counted with featureCounts v2.0.3 (60) and normalized/analyzed with DESeq2 v1.42.1 (61). DEGs (adj. p < 0.05, Benjamini-Hochberg correction) were annotated via VectorBase. GO-terms were simplified using the R package *simplifyEnrichment* v1.12.0. The KEGG database was used for pathway annotation.

### *De novo* transcriptome analysis *As. taiwanensis*

Unmapped reads were extracted with Samtools v1.15.1 (64) and assembled de novo following the recommendation of Bretaudeau *et al*. (62). Trinity v2.15.1 (63) was used with the Salmon method (64) for quantification. Transcripts were aligned to 17 Apicomplexan genomes from CryptoDB (https://cryptodb.org/cryptodb/app/, downloaded 12 Dec 2023). *Ascogregarina taiwanensis* is the only Apicomplexan species known to be part of the *Ae. albopictus* microbiota (18) and the CryptoDB database includes *Gregarinasina* as well as phylogenetically close taxa. A total of 4,676 transcripts were retained as *As. taiwanensis* and deposited at Zenodo (https://zenodo.org/records/14899302). DEG analysis followed the same pipeline as for mosquitoes, and annotations (GO-terms and KEGG pathways) were performed using CryptoDB.

### Quantification of dry mass, protein, uric acid and uricase activity

Ten pools of three females per condition were freeze-dried (8 h) and weighed. Samples were homogenized in 200 µL cold PBS (pH 7.4) with three 3 mm beads using a FastPrep-24 system (30 s, 6 m/s), centrifuged (1000 g, 5 min), and supernatants were transferred to black 1.5 mL microtubes (Heatrow Scientific). Protein content was measured using the Qubit Protein Assay Kit and a Qubit4 fluorometer (Invitrogen). Uric acid and uricase activity were quantified using the Amplex Red Kit (Invitrogen), with fluorescence measured at 570 nm (excitation at 530 nm) on a SpectraMAX iD3 plate reader (Molecular Devices). Data were converted using standard curves.

### Statistical analysis

All analyses were conducted in R v4.4.0 (65). Female ID was used as a random effect in mixed models. Parasite prevalence over time and larval prevalence were analyzed via binomial GLMMs using the glmmTMB package. Abdomen width data were transformed (Box-Cox) and analyzed using linear models (*stats, MASS*). Egg chamber, yolk area, and larval size were assessed with linear mixed models (*lme4*). Egg-laying dynamics were modeled with Poisson GLMMs (*glmmTMB*). Hatching rate was analyzed via binomial GLMMs (*lme4*). Protein, uric acid, and uricase data were normalized to dry mass and analyzed using linear mixed-effects models (*lme4*). Significance was tested using type II ANOVA or Wald Chi-square tests (*car*), followed by Tukey-HSD *post hoc* tests (*emmeans*). PCA and RDA (with permutation tests) of transcriptomic data were performed using *DESeq2* and *vegan*. Datasets are available at Zenodo (https://zenodo.org/records/14899302).

## Acknowledgments

This study was performed within the framework of the EC2CO CNRS project (year: 2020, name: Interasco, holder: Guillaume Minard) and the ANR JCJC project (year: 2024, name: Evasion, holder: Guillaume Minard). The Ph. D. of Maxime Girard was supported by a French ministerial fellowship delivered by the Ecology, Evolution, Microbiology and Modeling doctoral school of the University Claude Bernard Lyon 1. We thank the Master of Microbiology from the University Claude Bernard Lyon 1 and the Master of Microbiology from the University of Clermont Auvergne in which Mathieu Lays, Melanie Bretton and An-nah Chanfi were involved during their traineeship. Since none of the authors are native English speakers, the final draft of this manuscript was reviewed by an artificial intelligence software to improve its clarity. The authors carefully reviewed the suggested modifications and checked that it did not affect its meaning.

