## Supporting information for "Friend or foe: A mosquito parasite with mixed transmission mode displays mutualistic traits promoting oogenesis"

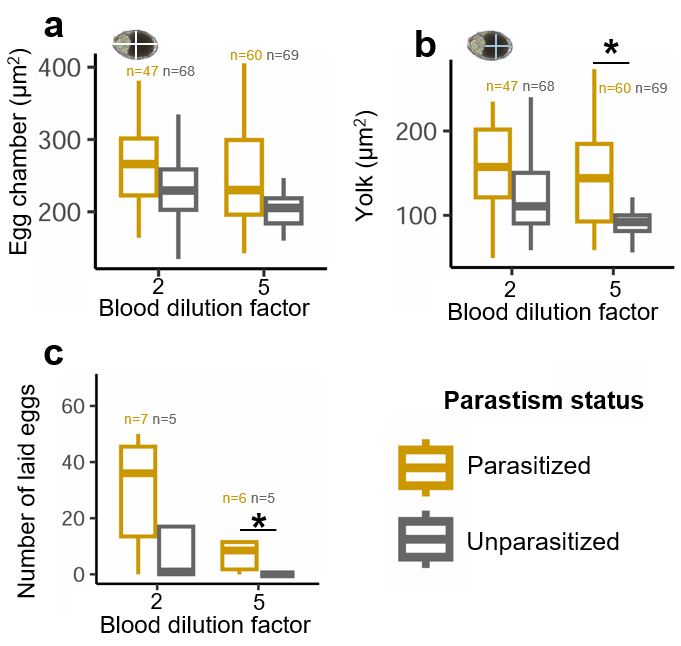

**Figure S1. Impact of *As. taiwanensis* and blood quality in oogenesis and egg laying.** (a) The follicle primary chamber and (b) yolk areas are reported for parasitized and unparasitized females 1DABM female mosquitoes using 1/2 or 1/5 diluted blood. (c) The number of eggs laid in each of those conditions was reported. Asterisks represent significant pairwise differences at a threshold of p ≤ 0.05 from (b) post hoc Tukey HSD and (c) Wilcoxon rank-sum tests.

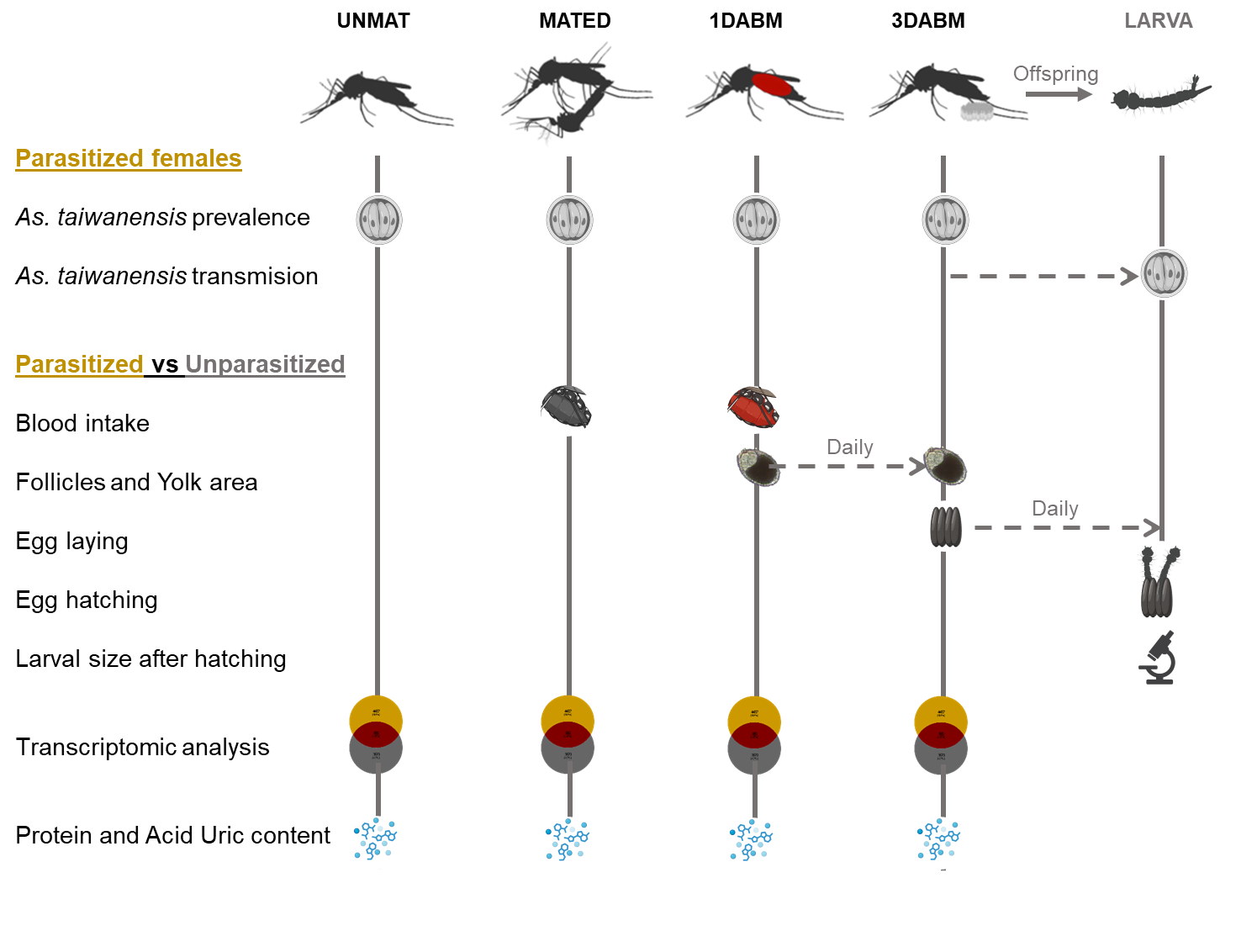
**Figure S1. Overview of the experimental design**. *Ascogregarina taiwanensis* prevalence was assessed in *Ae. albopictus* females at four life stages: unmated (UNMAT), mated (MATED), one day after blood meal (1DABM), and three days after blood meal (3DABM). Parasite transmission to offspring was evaluated via water and egg smearing. Blood intake was quantified by measuring abdomen width before and after feeding. Oogenesis was monitored by tracking primary follicle and yolk development over time. Egg-laying dynamics and larval size were recorded. Comparative transcriptomic analyses were performed to assess both parasite gene expression and its impact on the mosquito transcriptome. Protein content, uric acid levels, and uricase activity were also measured at each stage.

**Table S1. Metabolic pathways identified with the DEGs of 1 one day after blood meal (1DABM) parasitized females**

| **Condition** | **Parasitism status** | **KEGG_ID** | **Pathways** | **Fold Enrichment** |
| --- | --- | --- | --- | --- |
| UNMAT | Unparasitized | ec00983 | Drug metabolism - other enzymes | 1.63 |
|  |  | ec00332 | Carbapenem biosynthesis | 1.57 |
|  |  | ec00790 | Folate biosynthesis | 1.66 |
|  |  | ec00627 | Aminobenzoate degradation | 1.48 |
|  |  | ec00541 | O-Antigen nucleotide sugar biosynthesis | 1.49 |
|  |  | ec00500 | Starch and sucrose metabolism | 1.43 |
|  |  | ec00643 | Styrene degradation | 1.77 |
|  |  | ec00980 | Metabolism of xenobiotics by cytochrome P450 | 1.46 |
|  |  | ec00361 | Chlorocyclohexane and chlorobenzene degradation | 1.54 |
|  | Parasitized | ec00620 | Pyruvate metabolism | 1.96 |
|  |  | ec00220 | Arginine biosynthesis | 2.08 |
| MATED | Unparasitized | ec00071 | Fatty acid degradation | 2.39 |
|  |  | ec00901 | Indole alkaloid biosynthesis | 2.47 |
|  |  | ec00830 | Retinol metabolism | 1.99 |
|  |  | ec00905 | Brassinosteroid biosynthesis | 3.02 |
|  |  | ec00232 | Caffeine metabolism | 2.99 |
|  |  | ec00643 | Styrene degradation | 2.84 |
|  |  | ec00943 | Isoflavonoid biosynthesis | 3.02 |
|  |  | ec00522 | Biosynthesis of 12-, 14- and 16-membered macrolides | 2.25 |
|  |  | ec00982 | Drug metabolism - cytochrome P450 | 1.95 |
|  |  | ec00363 | Bisphenol degradation | 1.95 |
|  |  | ec00120 | Primary bile acid biosynthesis | 2.32 |
|  |  | ec00941 | Flavonoid biosynthesis | 2.15 |
|  |  | ec00361 | Chlorocyclohexane and chlorobenzene degradation | 2.28 |
|  |  | ec00980 | Metabolism of xenobiotics by cytochrome P450 | 1.96 |
|  |  | ec00904 | Diterpenoid biosynthesis | 2.21 |
|  |  | ec00903 | Limonene and pinene degradation | 1.96 |
|  |  | ec00906 | Carotenoid biosynthesis | 1.93 |
|  |  | ec00590 | Arachidonic acid metabolism | 1.84 |
|  |  | ec00591 | Linoleic acid metabolism | 1.94 |
|  |  | ec00260 | Glycine, serine and threonine metabolism | 2.05 |
|  |  | ec00140 | Steroid hormone biosynthesis | 1.79 |
|  |  | ec00945 | Stilbenoid, diarylheptanoid and gingerol biosynthesis | 1.76 |
|  |  | ec00966 | Glucosinolate biosynthesis | 1.85 |
|  |  | ec00623 | Toluene degradation | 1.77 |
|  |  | ec00626 | Naphthalene degradation | 1.84 |
|  |  | ec00625 | Chloroalkane and chloroalkene degradation | 2.08 |
|  |  | ec00380 | Tryptophan metabolism | 1.54 |
|  |  | ec01057 | Biosynthesis of type II polyketide products | 1.6 |
|  |  | ec00750 | Vitamin B6 metabolism | 3.22 |
|  |  | ec00051 | Fructose and mannose metabolism | 1.55 |
|  |  | ec00627 | Aminobenzoate degradation | 1.55 |
|  |  | ec00404 | Staurosporine biosynthesis | 1.68 |
|  |  | ec00500 | Starch and sucrose metabolism | 1.53 |
|  |  | ec00360 | Phenylalanine metabolism | 1.63 |
|  |  | ec00981 | Insect hormone biosynthesis | 1.79 |
|  |  | ec00130 | Ubiquinone and other terpenoid-quinone biosynthesis | 1.57 |
|  | Parasitized | ec00521 | Streptomycin biosynthesis | 3.32 |
|  |  | ec00410 | beta-Alanine metabolism | 2.68 |
|  |  | ec00052 | Galactose metabolism | 1.88 |
|  |  | ec00513 | Various types of N-glycan biosynthesis | 2.38 |
|  |  | ec00600 | Sphingolipid metabolism | 2.04 |
| 1DABM | Unparasitized | ec00521 | Streptomycin biosynthesis | 2.03 |
|  |  | ec00563 | Glycosylphosphatidylinositol (GPI)-anchor biosynthesis | 1.62 |
|  | Parasitized | ec00470 | D-Amino acid metabolism | 1.96 |
|  |  | ec00480 | Glutathione metabolism | 1.89 |
|  |  | ec00982 | Cytochrome P450 metabolism | 1.7 |
|  |  | ec00310 | Lysine degradation | 1.61 |
|  |  | ec00410 | beta-Alanine metabolism | 1.58 |
|  |  | ec00071 | Fatty acid degradation | 1.51 |
|  |  | ec00232 | Caffeine metabolism | 1.49 |
|  |  | ec00140 | Steroid hormone biosynthesis | 1.47 |
|  |  | ec00053 | Ascorbate and aldarate metabolism | 1.43 |
|  |  | ec00620 | Pyruvate metabolism | 1.43 |
| 3DABM | Unparasitized | ec00830 | Retinol metabolism | 2.07 |
|  |  | ec00901 | Indole alkaloid biosynthesis | 2.5 |
|  |  | ec00980 | Metabolism of xenobiotics by cytochrome P450 | 2.23 |
|  |  | ec00982 | Drug metabolism - cytochrome P450 | 2.07 |
|  |  | ec00591 | Linoleic acid metabolism | 2.26 |
|  |  | ec00232 | Caffeine metabolism | 2.91 |
|  |  | ec00363 | Bisphenol degradation | 2.03 |
|  |  | ec00120 | Primary bile acid biosynthesis | 2.45 |
|  |  | ec00140 | Steroid hormone biosynthesis | 2.1 |
|  |  | ec00905 | Brassinosteroid biosynthesis | 2.84 |
|  |  | ec00943 | Isoflavonoid biosynthesis | 2.83 |
|  |  | ec00590 | Arachidonic acid metabolism | 1.94 |
|  |  | ec00643 | Styrene degradation | 2.5 |
|  |  | ec00071 | Fatty acid degradation | 1.95 |
|  |  | ec00623 | Toluene degradation | 1.92 |
|  |  | ec00903 | Limonene and pinene degradation | 1.91 |
|  |  | ec00626 | Naphthalene degradation | 1.88 |
|  |  | ec00625 | Chloroalkane and chloroalkene degradation | 2.09 |
|  |  | ec00130 | Ubiquinone and other terpenoid-quinone biosynthesis | 1.79 |
|  |  | ec00361 | Chlorocyclohexane and chlorobenzene degradation | 1.97 |
|  |  | ec00261 | Monobactam biosynthesis | 1.73 |
|  |  | ec00966 | Glucosinolate biosynthesis | 1.8 |
|  |  | ec00904 | Diterpenoid biosynthesis | 1.92 |
|  |  | ec00053 | Ascorbate and aldarate metabolism | 1.59 |
|  |  | ec00940 | Phenylpropanoid biosynthesis | 1.59 |
|  |  | ec00500 | Starch and sucrose metabolism | 1.55 |
|  |  | ec00941 | Flavonoid biosynthesis | 1.75 |
|  |  | ec00260 | Glycine, serine and threonine metabolism | 1.81 |
|  |  | ec00522 | Biosynthesis of 12-, 14- and 16-membered macrolides | 1.72 |
|  |  | ec00983 | Drug metabolism - other enzymes | 1.43 |
|  |  | ec00906 | Carotenoid biosynthesis | 1.58 |
|  |  | ec00365 | Furfural degradation | 1.72 |
|  |  | ec00290 | Valine, leucine and isoleucine biosynthesis | 3.01 |
|  | Parasitized | ec00730 | Thiamine metabolism | 1.81 |
|  |  | ec00532 | Glycosaminoglycan biosynthesis - chondroitin sulfate / dermatan sulfate | 5.4 |
|  |  | ec00622 | Xylene degradation | 3.81 |
|  |  | ec00621 | Dioxin degradation | 3.6 |
|  |  | ec00260 | Glycine, serine and threonine metabolism | 1.99 |
|  |  | ec00770 | Pantothenate and CoA biosynthesis | 2.11 |

**Table S2. Metabolic pathways identified in the *As. taiwanensis* transcriptome at different stages of the female life cycle, *i.e.* unmated (UNMAT), mated (MATED), one day after blood meal (1DABM) and three days after blood meal (3DABM).**

| **Stages** | **KEGG_ID** | **Metabolic pathways** | **Fold Enrichment** |
| --- | --- | --- | --- |
| UNMAT | ec00280 | Valine, leucine and isoleucine degradation | 7.04 |
|  | ec00020 | Citrate cycle (TCA cycle) | 6.77 |
|  | ec00970 | Aminoacyl-tRNA biosynthesis | 5.46 |
|  | ec00260 | Glycine, serine and threonine metabolism | 5.03 |
|  | ec00640 | Propanoate metabolism | 5.03 |
| MATED | ec00401 | Novobiocin biosynthesis | 25.09 |
|  | ec00660 | C5-Branched dibasic acid metabolism | 12.55 |
|  | ec00400 | Phenylalanine, tyrosine and tryptophan biosynthesis | 5.46 |
|  | ec00020 | Citrate cycle (TCA cycle) | 5.25 |
|  | ec00620 | Pyruvate metabolism | 3.33 |
| 1DABM | ec00401 | Novobiocin biosynthesis | 28.68 |
|  | ec00660 | C5-Branched dibasic acid metabolism | 19.85 |
|  | ec00220 | Arginine biosynthesis | 8.19 |
|  | ec00250 | Alanine, aspartate and glutamate metabolism | 7.44 |
|  | ec00020 | Citrate cycle (TCA cycle) | 6.87 |
| 3DABM | ec00430 | Taurine and hypotaurine metabolism | 10.75 |
|  | ec00220 | Arginine biosynthesis | 5.46 |
|  | ec00020 | Citrate cycle (TCA cycle) | 4.96 |
|  | ec00730 | Thiamine metabolism | 4.61 |
|  | ec00230 | Purine metabolism | 4.21 |
